# Cerebellar and subcortical atrophy contribute to psychiatric symptoms in frontotemporal dementia

**DOI:** 10.1101/2021.11.12.468429

**Authors:** Aurélie Bussy, Jake Levy, Tristin Best, Raihaan Patel, Lani Cupo, Tim Van Langenhove, Jorgen Nielsen, Yolande Pijnenburg, Maria Landqvist Waldö, Anne Remes, Matthias L Schroeter, Isabel Santana, Florence Pasquier, Markus Otto, Adrian Danek, Johannes Levin, Isabelle Le Ber, Rik Vandenberghe, Matthis Synofzik, Fermin Moreno, Alexandre de Mendonça, Raquel Sanchez-Valle, Robert Laforce, Tobias Langheinrich, Alexander Gerhard, Caroline Graff, Chris R. Butler, Sandro Sorbi, Lize Jiskoot, Harro Seelaar, John C. van Swieten, Elizabeth Finger, Maria Carmela Tartaglia, Mario Masellis, Pietro Tiraboschi, Daniela Galimberti, Barbara Borroni, James B. Rowe, Martina Bocchetta, Jonathan D. Rohrer, Gabriel A. Devenyi, M. Mallar Chakravarty, Simon Ducharme, on behalf of the GENetic Frontotemporal dementia Initiative (GENFI)

**Affiliations:** Computional Brain Anatomy (CoBrA) Laboratory, Cerebral Imaging Centre, Douglas Mental Health University Institute, Montreal, Quebec, Canada; Integrated Program in Neuroscience, McGill University, Montreal, Canada; Department of Biomedical Engineering, McGill University, Montreal, Quebec, Canada; Department of Molecular Genetics, VIB, Antwerp, Belgium; Institute Born-Bunge, University of Antwerp, Antwerp, Belgium; Department of Neurology, Danish Dementia Research Centre, Rigshospitalet, University of Copenhagen, Denmark; Section of Neurogenetics, Department of Cellular and Molecular Medicine, University of Copenhagen, Copenhagen, Denmark; Alzheimer Center Amsterdam, Department of Neurology, Amsterdam Neuroscience, Vrije Universiteit Amsterdam, Amsterdam UMC, Amsterdam, The Netherlands; Division of Clinical Sciences Helsingborg, Department of Clinical Sciences Lund, Lund University, Lund, Sweden; Unit of Clinical Neuroscience, Neurology, University of Oulu, Oulu, Finland; Medical Research Center, Oulu University Hospital, Oulu, Finland; Max Planck Institute for Human Cognitive and Brain Sciences, Leipzig; Clinic for Cognitive Neurology, University Clinic Leipzig, Leipzig, Germany; University Hospital of Coimbra (HUC), Neurology Service, Faculty of Medicine, University of Coimbra, Coimbra, Portugal; Center for Neuroscience and Cell Biology, Faculty of Medicine, University of Coimbra, Coimbra, Portugal; Universite de Lille, France; Inserm 1172, Lille, France; CHU, CNR-MAJ, Labex Distalz, LiCEND Lille, France; Department of Neurology, University of Ulm, Ulm, Germany; Neurologische Klinik und Poliklinik, Ludwig-Maximilians-Universität, Munich; German Center for Neurodegenerative Diseases (DZNE); Munich Cluster of Systems Neurology, Munich, Germany; Sorbonne Université, Paris Brain Institute – Institut du Cerveau – ICM, Inserm U1127, CNRS UMR 7225, AP-HP - Hôpital Pitié-Salpêtrière, Paris, France; Centre de référence des démences rares ou précoces, IM2A, Département de Neurologie, AP-HP - Hôpital Pitié-Salpêtrière, Paris, France; Département de Neurologie, AP-HP - Hôpital Pitié-Salpêtrière, Paris, France; Laboratory for Cognitive Neurology, Department of Neurosciences, KU Leuven, Leuven, Belgium; Neurology Service, University Hospitals Leuven, Belgium; Leuven Brain Institute, KU Leuven, Leuven, Belgium; Department of Neurodegenerative Diseases, Hertie-Institute for Clinical Brain Research and Center of Neurology, University of Tübingen, Tübingen, Germany; Neuroscience Area, Biodonostia Health Research Institute, San Sebastian, Gipuzkoa, Spain; Cognitive Disorders Unit, Department of Neurology, Donostia University Hospital, San Sebastian, Gipuzkoa, Spain; Faculty of Medicine, University of Lisbon, Lisbon, Portugal; Alzheimer’s disease and Other Cognitive Disorders Unit, Neurology Service, Hospital Clínic, Institut d’Investigacións Biomèdiques August Pi I Sunyer, University of Barcelona, Barcelona, Spain; Clinique Interdisciplinaire de Mémoire, Département des Sciences Neurologiques, CHU de Québec, and Faculté de Médecine, Université Laval, QC, Canada; Division of Neuroscience and Experimental Psychology, Wolfson Molecular Imaging Centre, University of Manchester, Manchester, UK; Cerebral Function Unit, Manchester Centre for Clinical Neurosciences, Salford Royal NHS Foundation Trust, Salford, UK; Departments of Geriatric Medicine and Nuclear Medicine, University of Duisburg-Essen, Germany; Center for Alzheimer Research, Division of Neurogeriatrics, Department of Neurobiology, Care Sciences and Society, Bioclinicum, Karolinska Institutet, Solna, Sweden; Unit for Hereditary Dementias, Theme Aging, Karolinska University Hospital, Solna, Sweden; Nuffield Department of Clinical Neurosciences, Medical Sciences Division, University of Oxford, Oxford, UK; Department of Brain Sciences, Imperial College London, UK; Department of Neurofarba, University of Florence, Italy; IRCCS Fondazione Don Carlo Gnocchi, Florence, Italy; Department of Neurology, Erasmus Medical Centre, Rotterdam, Netherlands; Department of Clinical Neurological Sciences, University of Western Ontario, London, Ontario Canada; Tanz Centre for Research in Neurodegenerative Diseases, University of Toronto, Toronto, Canada; Sunnybrook Health Sciences Centre, Sunnybrook Research Institute, University of Toronto, Toronto, Canada; Fondazione IRCCS Istituto Neurologico Carlo Besta, Milano, Italy; Department of Biomedical, Surgical and Dental Sciences, University of Milan, Milan, Italy; Fondazione IRCCS Ca’ Granda, Ospedale Maggiore Policlinico, Milan, Italy; Centre for Neurodegenerative Disorders, Department of Clinical and Experimental Sciences, University of Brescia, Brescia, Italy; Department of Clinical Neurosciences and Cambridge University Hospitals NHS Trust and Medical Research Council Cognition and Brain Sciences Unit, University of Cambridge, Cambridge, UK; Department of Neurodegenerative Disease, Dementia Research Centre, UCL Institute of Neurology, Queen Square, London, UK; Department of Psychiatry, McGill University, Montreal, Quebec, Canada; Douglas Mental Health University Institute, Department of Psychiatry, McGill University, Montreal, Quebec, Canada; McConnell Brain Imaging Centre, Montreal Neurological Institute, McGill University, Montreal, Quebec, Canada

**Keywords:** frontotemporal dementia, mutation, morphometry, neuropsychiatric symptoms, cerebellum

## Abstract

Recent studies have suggested that cerebellar and subcortical structures are impacted early in the disease progression of genetic frontotemporal dementia (FTD) due to microtubule-associated protein tau (*MAPT*), progranulin (*GRN*) and chromosome 9 open reading frame 72 (*C9orf72*). However, the clinical contribution of the structures involved in the cerebello-subcortical circuitry has been understudied in FTD given their potentially central role in cognition and behaviour processes. The present study aims to investigate whether there is an association between the atrophy of the cerebellar and subcortical structures, and neuropsychiatric symptoms (using the revised version of the Cambridge Behavioral Inventory, CBI-R) across genetic mutations and whether this association starts during the preclinical phase of the disease. Our study included 983 participants from the Genetic Frontotemporal dementia Initiative (GENFI) including mutation carriers (n=608) and non-carrier first-degree relatives of known symptomatic carriers (n= 375). Voxel-wise analysis of the thalamus, striatum, globus pallidus, amygdala, and the cerebellum was performed using deformation based morphometry (DBM) and partial least squares analyses (PLS) were used to link morphometry and behavioural symptoms. Our univariate results suggest that in this group of primarily presymptomatic subjects, volume loss in subcortical and cerebellar structure was primarily a function of aging, with only the *C9orf72* group showing more pronounced volume loss in the thalamus compared to the non-carrier individuals. PLS analyses demonstrated that the cerebello-subcortical circuitry is related to all neuropsychiatric symptoms from the CBI-R, with significant overlap in brain/behaviour patterns, but also specificity for each genetic group. The biggest differences were in the extent of the cerebellar involvement (larger extent in *C9orf72* group) and more prominent amygdalar contribution in the *MAPT* group. Finally, our findings demonstrated that *C9orf72* and *MAPT* brain scores were related to estimated years before the age of symptom onset (EYO) in a second order relationship highlighting a steeper brain score decline 20 years before expected symptom onset, while *GRN* brain scores were related to age and not EYO. Overall, these results demonstrated the important role of the subcortical structures and especially of the cerebellum in genetic FTD symptom expression.

## 1. Introduction

Frontotemporal dementia (FTD) is the second most common form of neurodegenerative dementia in people under 65 years of age ^1,2^. Behavioural and personality alterations encompassing ‘negative’ symptoms (apathy, loss of empathy) and ‘positive’ symptoms (disinhibition, inappropriate social behaviour) are among the most prominent symptoms of FTD ^3,4^. While most FTD cases are sporadic, 10-20% of cases are caused by three well known autosomal dominant full penetrance mutations: microtubule-associated protein tau (*MAPT*), progranulin (*GRN*) and chromosome 9 open reading frame 72 (*C9orf72*) mutations ^5–7^. Studying presymptomatic mutation carriers can provide valuable insight into the neuroanatomical modifications that occur during the preclinical phase of FTD. Indeed, there is mounting evidence demonstrating that FTD-related pathophysiology starts several years before the clear onset of the disease ^8–10^. Furthermore, psychosis-related symptoms can constitute the prodrome of genetic FTD ^11^, and emerging evidence suggests that presymptomatic carriers have a subtle increase in neuropsychiatric symptoms ^9,12^.

To date, most FTD studies have predominantly tried to relate cognitive and behavioural impairment with cortical atrophy ^10,13–17^. However, there is a growing number of papers demonstrating that the atrophy of the cerebellum and subcortical structures in genetic FTD (particularly in the *C9orf72* expansion carriers) appears earlier than the cortical impairment in the disease progression. Indeed, a previous study has shown that presymptomatic FTD mutation carriers demonstrate atrophy in subcortical regions such as in the thalamus, striatum and amygdala, as well as in the cerebellum, preceding larger cortical atrophy, compared to non-carriers individuals ^18^. Moreover, we know from numerous studies in healthy controls that these structures have been characterized for their central role in various cognitive and behavioural processes such as arousal, attention, mood, motivation, language, memory, abstraction, and visuospatial skills ^19–23^. Therefore, we could expect that a disruption of these structures in FTD could affect many of these cognitive and behavioural functions. Indeed, a recent study examined the distinct relationships between subcortical structures and neuropsychiatric symptoms (hallucinations, delusions, depression and anxiety) in the three main forms of genetic FTD and demonstrated that cerebellar atrophy was correlated with anxiety in *C9orf72* carriers, further implicating the cerebellar structure in FTD symptomatology ^24^. Therefore, the goal of this study is to further investigate whether there is an association between the atrophy of the regions of the cerebello-subcortical circuitry and diverse FTD-specific cognitive and behavioural metrics across genetic mutations and whether this association starts during the preclinical phase of the disease.

The present study used a dataset gathering 983 participants from the Genetic Frontotemporal dementia Initiative [GENFI; ^9^], a unique sample given that most FTD studies ^25,26^ have often been limited by a small number of individuals (<100). Here we focus specifically on a voxel-wise analysis in a region of interest encompassing the thalamus, striatum, globus pallidus, amygdala, and the cerebellum using deformation-based morphometry (DBM). Of note, these regions have been selected because of their implication in the cerebello-subcortical circuitry and for their potential implication in psychiatric symptoms. We first examined the relationship between voxel-wise measures and FTD-relevant demographic and clinical data, such as: genetic mutation status, age, estimated years before the age of symptom onset (EYO; calculated as the difference between the parental age of symptom onset and the individual’s current age ^27^) and symptomatic status (symptomatic or non-symptomatic). Next, a multivariate technique (Partial least squares analyses [PLS]) was used to derive linked dimensions of voxel-wise morphometry with cognitive and behavioural symptoms and extract different patterns specific to each mutation group. We finally investigated if these brain/behavioural patterns could be explained by different demographic and clinical information listed previously.

## 2. Methods

### 2.1. Participants

983 participants were selected from the GENFI2 dataset data release 5 which includes data from participants across multiple research sites in the UK, the Netherlands, Belgium, France, Spain, Portugal, Italy, Germany, Sweden, Finland and Canada. The participants were either known carriers of a pathogenic mutation in *MAPT* (n= 104), *GRN* (n=243), *C9orf72* (n= 256), *TANK-binding kinase 1* (*TBK1*; n=5) or first-degree relatives of known symptomatic carriers (“non-carriers” group; n= 375). *TBK1* carriers were excluded because of the low number of carriers of this mutation. Details on the participants selected for further analyses (see 2.3.2 Raw quality control, 2.3.3 Preprocessing sections and 3.1. Demographic and clinical information) can be found in Table 1.

**Table 1.** Complete demographic and clinical information of the 730 individuals who passed motion QC.

|  | Non carriers<br>(n=281) | *C9orf72* carriers<br>(n=184) | *GRN* carriers<br>(n=185) | *MAPT* carriers<br>(n=80) | Total<br>(n=730) |
| --- | --- | --- | --- | --- | --- |
| **Age at visit**<br>Mean (SD)<br>Median [Min, Max] | 46.1 (13.5)<br>44.6 [19.4, 85.0] | 50.6 (14.1)<br>51.4 [20.1, 77.9] | 50.5 (13.3)<br>51.5 [20.2, 77.0] | 44.8 (13.9)<br>43.4 [20.5, 74.0] | 48.2 (13.8)<br>47.5 [19.4, 85.0] |
| **EYO**<br>Mean (SD)<br>Median [Min, Max] | -13.1 (14.1)<br>-15.3 [-50.0,27.6] | -8.0 (14.0)<br>-6.53 [-47.4, 1.3] | -10.1 (13.2)<br>-8.83 [-39.0, 19.1] | -7.94 (13.7)<br>-6.82 [-39.4, 15.7] | -10.6 (13.9)<br>-11.4 [-50.0,27.6] |
| **Gender**<br>Female<br>Male | 164 (58.4%)<br>117 (41.6%) | 96 (52.2%)<br>88 (47.8%) | 119 (64.3%)<br>66 (35.7%) | 40 (50.0%)<br>40 (50.0%) | 419 (57.4%)<br>311 (42.6%) |
| **Genetic status**<br>Non-symptomatic<br>Carrier presymptomatic<br>Carrier symptomatic | 281 (100%)<br>0 (0%)<br>0 (0%) | 0 (0%)<br>122 (66.3%)<br>62 (33.7%) | 0 (0%)<br>137 (74.1%)<br>48 (26.0 %) | 0 (0%)<br>56 (70.0%)<br>24 (30.0 %) | 281 (38.5%)<br>315 (43.1%)<br>134 (18.4%) |
| **Memory**<br>Mean (SD)<br>Median [Min, Max] | 1.1 (2.15)<br>0.0 [0.0, 15.0] | 5.9 (7.8)<br>2.0 [0.0, 28.0] | 3.6 (6.2)<br>1.0 [0.0, 32.0] | 5.0 (7.6)<br>1.0 [0.0, 30.0] | 3.42 (6.1)<br>0.5 [0.0, 32.0] |
| **Everyday skills**<br>Mean (SD)<br>Median [Min, Max] | 0.1 (0.7)<br>0.0 [0.0, 9.0] | 2.8 (5.0)<br>0.0 [0.0, 20.0] | 2.0 (4.8)<br>0.0 [0.0,20.0] | 1.5 (3.4)<br>0.0 [0.0, 14.0] | 1.4 (3.8)<br>0.0 [0.0, 20.0] |
| **Self care**<br>Mean (SD)<br>Median [Min, Max] | 0.0 (0.1)<br>0.0 [0.0, 1.0] | 1.5 (3.6)<br>0.0 [0.0, 16.0] | 1.0 (3.1)<br>0.0 [0.0, 16.0] | 0.5 (1.6)<br>0.0 [0.0, 8.0] | 0.7 (2.5)<br>0.0 [0.0, 16.0] |
| **Abnormal behaviour**<br>Mean (SD)<br>Median [Min, Max] | 0.5 (1.1)<br>0.0 [0.0, 7.0] | 3.2 (4.7)<br>1.0 [0.0, 19.0] | 1.7 (3.6)<br>0.0 [0.0, 23.0] | 2.8 (4.8)<br>0.5 [0.0, 20.0] | 1.8 (3.6)<br>0.0 [0.0, 23.0] |
| **Sleep**<br>Mean (SD)<br>Median [Min, Max] | 0.4 (1.0)<br>0.0 [0.0, 6.0] | 1.2 (1.9)<br>0.0 [0.0, 8.0] | 1.0 (1.7)<br>0.0 [0.0, 8.0] | 1.4 (1.9)<br>1.0 [0.0, 8.0] | 0.9 (1.6)<br>0.0 [0.0, 8.0] |
| **Eating**<br>Mean (SD)<br>Median [Min, Max] | 0.2 (0.9)<br>0.0 [0.0, 12.0] | 2.2 (4.1)<br>0.0 [0.0,16.0] | 1.3 (2.9)<br>0.0 [0.0, 14.0] | 2.0 (4.0)<br>0.0 [0.0, 15.0] | 1.2 (3.0)<br>0.0 [0.0, 16.0] |
| **Mood**<br>Mean (SD)<br>Median [Min, Max] | 0.9 (1.6)<br>0.0 [0.0, 9.0] | 2.2 (2.8)<br>1.0 [0.0, 16.0] | 1.5 (2.5)<br>0.0 [0.0, 12.0] | 2.4 (2.9)<br>1.5 [0.0, 13] | 1.6 (2.4)<br>0.0 [0.0, 16.0] |
| **Beliefs**<br>Mean (SD)<br>Median [Min, Max] | 0.0 (0.1)<br>0.0 [0.0, 1.0] | 0.4 (1.4)<br>0.0 [0.0, 12.0] | 0.2 (1.1)<br>0.0 [0.0, 10.0] | 0.1 (0.6)<br>0.0 [0.0, 4.0] | 0.2 (1.0)<br>0.0 [0.0, 12.0] |
| **Stereotypic behaviour**<br>Mean (SD)<br>Median [Min, Max] | 0.4 (1.2)<br>0.0 [0.0, 8.0] | 2.6 (3.8)<br>1.0 [0.0, 16.0] | 1.2 (2.4)<br>0.0 [0.0, 12.0] | 2.7 (4.7)<br>0.0 [0.0, 16.0] | 1.4 (3.0)<br>0.0 [0.0, 16.0] |
| **Motivation**<br>Mean (SD)<br>Median [Min, Max] | 0.5 (1.7)<br>0.0 [0.0, 16.0] | 3.9 (5.7)<br>0.0 [0.0, 20.0] | 2.3 (4.5)<br>0.0 [0.0, 20.0] | 3.0 (4.9)<br>0.0 [0.0, 19.0] | 2.1 (4.4)<br>0.0 [0.0, 20.0] |

### 2.2. Cognitive, behavioural and symptom assessments

All participants underwent clinical, cognitive and behavioural assessments. A “symptomatic status” binary variable was defined by clinicians (based on the clinician judgement at the time of the participant’s first GENFI visit), where the individuals were either assigned as “non-symptomatic” if they did not demonstrate overall symptoms, or “symptomatic” if they expressed FTD symptoms (this variable is not used in PLS analyses, but used in lmer and PLS post-hoc analyses; see 2.4.1. Linear mixed effect models and 2.4.3. Post-hoc analyses of PLS outputs).

Further, a validated scale for FTD behavioural assessment, the Cambridge Behavioural Inventory Revised version (CBI-R; ^28^), was performed to evaluate and quantify 10 specific cognitive, behavioural and affective symptoms as well as activities of daily living ^28^ and was used in PLS analyses (see 2.4.2. Partial least squares analysis). This assessment evaluates categories such as memory (memory, attention and orientation), everyday skills, self care, abnormal behaviour (challenging behaviour and disinhibition), mood (depression and agitation), beliefs (auditory and visual hallucinations; considered to be psychotic symptoms), eating, sleep, stereotypic behaviour (repetitive behaviour and motor movement) and motivation. The CBI-R test rates the frequency of any particular behaviour on a scale of 0-4. A score of 0 denotes no impairment, a score of 1 an occasional occurrence defined as a few times per month, 2 a repeated occurrence defined as a few times per week, 3 a daily occurrence, and 4 a constant occurrence. Scores of 3 or 4 are indicative of a severe behavioural deficit. The rating of each question is summed to give a final score per CBI-R category and these itemized scores were used in subsequent analyses; memory is rated on 32 points, abnormal behaviour on 24 points, everyday skills and motivation on 20 points, self care, mood, eating and stereotypic behaviour on 16 points, beliefs on 12 points and sleep on 8 points.

### 2.3 Image processing

#### 2.3.1. Image acquisition

All participants were recruited and scanned in a GENFI2 site. T1-weighted (T1w) images were acquired using an MPRAGE sequence (for parameters https://www.genfi.org/study/). Of the 730 participants included in the statistical analyses (see 2.3.2 Raw quality control, 2.3.3 Preprocessing sections and 3.1. Demographic and clinical information), 248 were scanned on a Philips 3T, 192 on a Siemens Trio 3T, 111 on a Siemens Skyra 3T, 32 on a Siemens 1.5T, 107 on a Siemens Prisma 3T, 3 on a Siemens 3T, 31 on a GE 3T and 6 on a GE 1.5T scanners. Scan protocols were designed at the outset of the study to ensure adequate matching between the scanners and image quality control.

#### 2.3.2. Raw quality control

Motion artifacts caused by involuntary movements such as cardiac, respiratory motion or drift over time are common in structural magnetic resonance (MR) images and negatively impact the quality of the data ^29–31^. In order to control for these artifacts, rigorous quality control (QC) of all raw images was performed (Figure 1) by two raters (JL, TB). The QC guidelines developed in the Computational Brain Anatomy (CoBrA) Laboratory (^32^; https://github.com/CoBrALab/documentation/wiki/Motion-Quality-Control-Manual) were followed and 130 scans were excluded due to motion artifacts.

**Figure 1:**
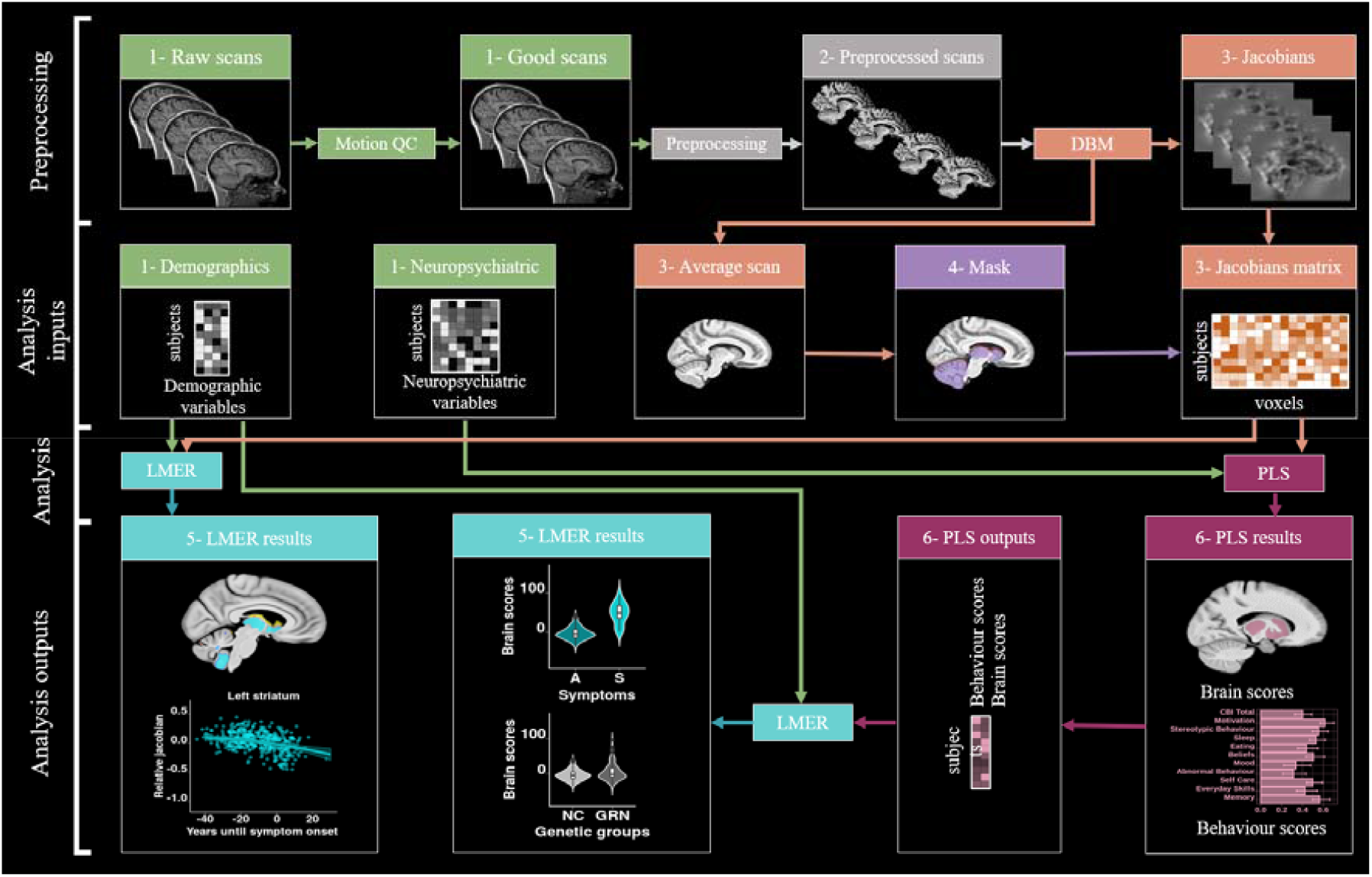
Chart flow of the step by step methods and analyses used in this paper. 1-Green: Raw inputs (2.3.2. Raw quality control); **2-Gray**: preprocessing (2.3.3 Preprocessing); **3-Orange**: Deformation based morphometry (2.3.4. Deformation based morphometry); **4-Purple**: Mask creation (2.3.5. Mask creation); **5-Turquoise**: linear mixed effect models (2.4.1. Linear mixed effect models); **6-Pink**: Partial least squares (2.4.2. Partial least squares analysis). Abbreviations: QC: quality control; Preprocessing: minc-bpipe-library; DBM: deformation based morphometry; SVD: singular value decomposition; PLS: partial least squares; LMER: linear mixed effect models.

#### 2.3.3. Preprocessing

Preprocessing steps were performed by two authors (JL, TB) on the raw T1 images in order to standardize images being input into the deformation-based analysis (see 2.3.4. Deformation based morphometry; Figure 1). The minc-bpipe-library pipeline (https://github.com/CobraLab/minc-bpipe-library) was used to perform the following steps: N4 bias field correction ^33^, registration to the Montreal Neurological Institute (MNI) space using bestlinreg ^34,35^, field-of-view standardization and brain orientation to MNI space using an inverse-affine transformation of a MNI space head mask, and brain extraction using BEaST technique ^36^. Additionally, minc-bpipe-library pipeline allows rapid supervision of these steps by providing quality control images of the various preprocessing stages.

#### 2.3.4. Deformation based morphometry

First, the skull-stripped preprocessed brains from minc-bpipe-library were used as inputs and were registered using a non-resampling rigid registration (6-parameters) to the MNI space. We used the two-level deformation based morphometry (DBM) python pipeline developed in the CoBrA Lab (https://github.com/CoBrALab/twolevel_ants_dbm) to investigate voxel-wise morphometry (Figure 1). Each individual image was warped using affine and non-linear registration to create an unbiased average using ANTs tools using a group-wise registration strategy ^37^. Relative voxel volume increases and decreases were determined from the deformation fields by estimating the Jacobian determinant at each voxel. This measure represents the relative difference at each voxel (as a proportion) relative to the group average (the residual affine components present in the nonlinear deformation field are also removed). This mathematical transformation allows easier statistical analyses and interpretation: positive values indicate that the voxel in template space must be expanded to get to the subject space, and negative values indicate that the voxel in template space must be reduced. The relative Jacobians were blurred with a 2 mm full-width-at-half-maximum 3D Gaussian to approximate the Gaussian assumptions required for the statistical field.

#### 2.3.5. Mask creation

In order to focus our voxel-wise analysis on the regions of interest (ROIs), a mask of the subcortical structures (striatum, globus pallidus, thalamus), the amygdala and the cerebellum was manually created on the average brain obtained from DBM analyses.

### 2.4. Statistics

#### 2.4.1. Linear mixed effect models

Vertex-wise linear mixed-effects models (*vertexLmer* from *RMINC_1*.*5*.*2*.*2* package in R 3.6.3) were used to test the significance of our relative Jacobians within our mask (Figure 1). This model included genetic mutation status (non-carriers, *C9orf72, GRN* or *MAPT* carriers), sex, education, symptomatic status (symptomatic/non-symptomatic), age at visit and EYO as fixed effects and, scanner and family ID as random effects (to control for potential relatives). Age and EYO were modeled as linear, second or third order using natural splines (ns from *splines* package) and Akaike information criterion [AIC; ^38^] was used to investigate the most appropriate relationships. The model with the lowest AIC was selected and was considered to best fit the data ^39^; see also previous work from our group ^32,40,41^. AIC results demonstrated that a second order relationship was the best fit for our analyses and therefore the model (eq. 1) was selected for the lmer analyses. An interaction between genetic mutation status and age was also tested in the model but was excluded due to high AIC values. A 5% false-discovery rate (FDR) correction was applied to control for the expected proportion of “discoveries” that are false ^42,43^.

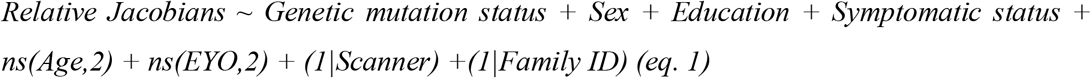

Here, given that the optimal model based on AIC excluded the interaction between genetic mutation status and age, we did not look at each mutation individually but rather we looked at the overall pattern of change with age and EYO. This approach is more stringent and gives us high confidence in the effects that we found, but consequently we do not test for relationships between specific mutations and age /EYO relationships. Of note, given our primary objective to look at mutation specific relationships between cerebral changes and psychiatric symptoms, our partial least squares analyses (2.4.2. Partial least squares) did test for each mutation brain and behaviour changes with age and EYO.

#### 2.4.2. Partial least squares

Partial least squares (PLS) correlation is a multivariate technique that is used to detect covariance patterns across two matrices via matrix decomposition techniques (Figure 1). The goal of PLS is to identify a set of latent variables (LVs) that explain patterns of covariance between brain data (here relative Jacobians) and behaviour data (here CBI-R scores) with the constraint that LVs explain as much of the covariance between the two matrices as possible. Theoretically, each LV depicts a linear combination of the brain and behaviour matrices. Here, four PLS analyses were run, one for each mutation group (*C9orf72, GRN, MAPT* and non-carriers separately), in order to be able to examine whether each mutation group would be associated with distinct brain/behaviour patterns. Our brain data included the relative Jacobian of each voxel for each subject (matrix size 3609356×184 for *C9orf72*; 3609356×185 for *GRN*, 3609356×80 for *MAPT* and 3609356×281 for non-carriers). Our behaviour data contained 10 CBI-R scores for each subject (matrix size 10×184 for *C9orf72*; 10×185 for *GRN*, 10×80 for *MAPT* and 10×281 for non-carriers). Note that this matrix does not contain any information on symptomatic vs presymptomatic, age or EYO.

Each LV was tested statistically using permutation testing following a similar protocol (detailed in the Supplementary methods) as in previous studies ^44–49^. Secondly, the degree to which each brain and behaviour variable contributes to these LVs was tested using a bootstrap resampling technique detailed in the Supplementary methods (BSR threshold of 2.58 = p-value of 0.01; Krishnan et al., 2011; McIntosh and Lobaugh, 2004; Nordin et al., 2018; Persson et al., 2014; Zeighami et al., 2017). Supplementary figure 1 displays the PLS brain scores using a less stringent threshold (BSR threshold of 1.96 = p-value of 0.05).

#### 2.4.3. Post-hoc analyses of PLS outputs

Once the PLS results were obtained and tested for significance, the brain scores and behaviour scores were further analyzed to determine if they were associated with key demographic and clinical variables. Linear mixed effect models including age, EYO, sex, education and symptomatic status as fixed effects and scanner and family ID as random effects to examine the brain scores (eq. 2) or the behaviour scores (eq. 3). False discovery rate (FDR) correction was applied to correct for multiple comparisons. We corrected for the eight p-values (one per predictor) x six models (one brain score (2) and one behaviour score (3) models per mutation group (*C9orf72, GRN* and *MAPT*). The goal of these analyses is to see if the LVs found from PLS capture patterns of brain and behaviour which are specific to disease-related demographics and symptomatic status. Indeed, PLS was blind to the EYO, sex, education and to the fact that some individuals were presymptomatic or symptomatic. These models were also tested on the presymptomatic carriers only (therefore without the “symptomatic status” variable). These results can be found in the supplementary figure 2.

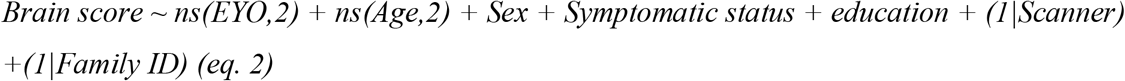

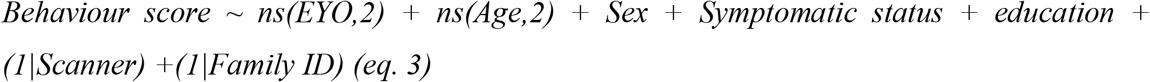

#### 2.4.4. Age and EYO model visualization

To visualize the significant age effects (*effects* package in R 3.6.3), we used the model coefficients to create the predicted Jacobians (in Figure 2) or the brain/behaviour scores (in Figure 4) every one year between age 19 and 85 for a subject of mean EYO with symptomatic status, sex and education as fixed effects and scanner and family ID as random effects. Inversely, to plot significant EYO effects, we used the model coefficients to create the predicted Jacobians (in Figure 2) or the brain/behaviour scores (in Figure 4) every one year between EYO −50 and 30 for a subject of mean age with symptomatic status, sex and education as fixed effects and scanner and family ID as random effects. The models were therefore computed using unweighted averages over the levels of non-focal factors such as sex, education, genetic mutation status and symptomatic status (See code example in the Supplementary methods). The advantage of this visualization process is to properly visualize the “significant” effects found in a model when there are multiple covariates also included in the model.

**Figure 2:**
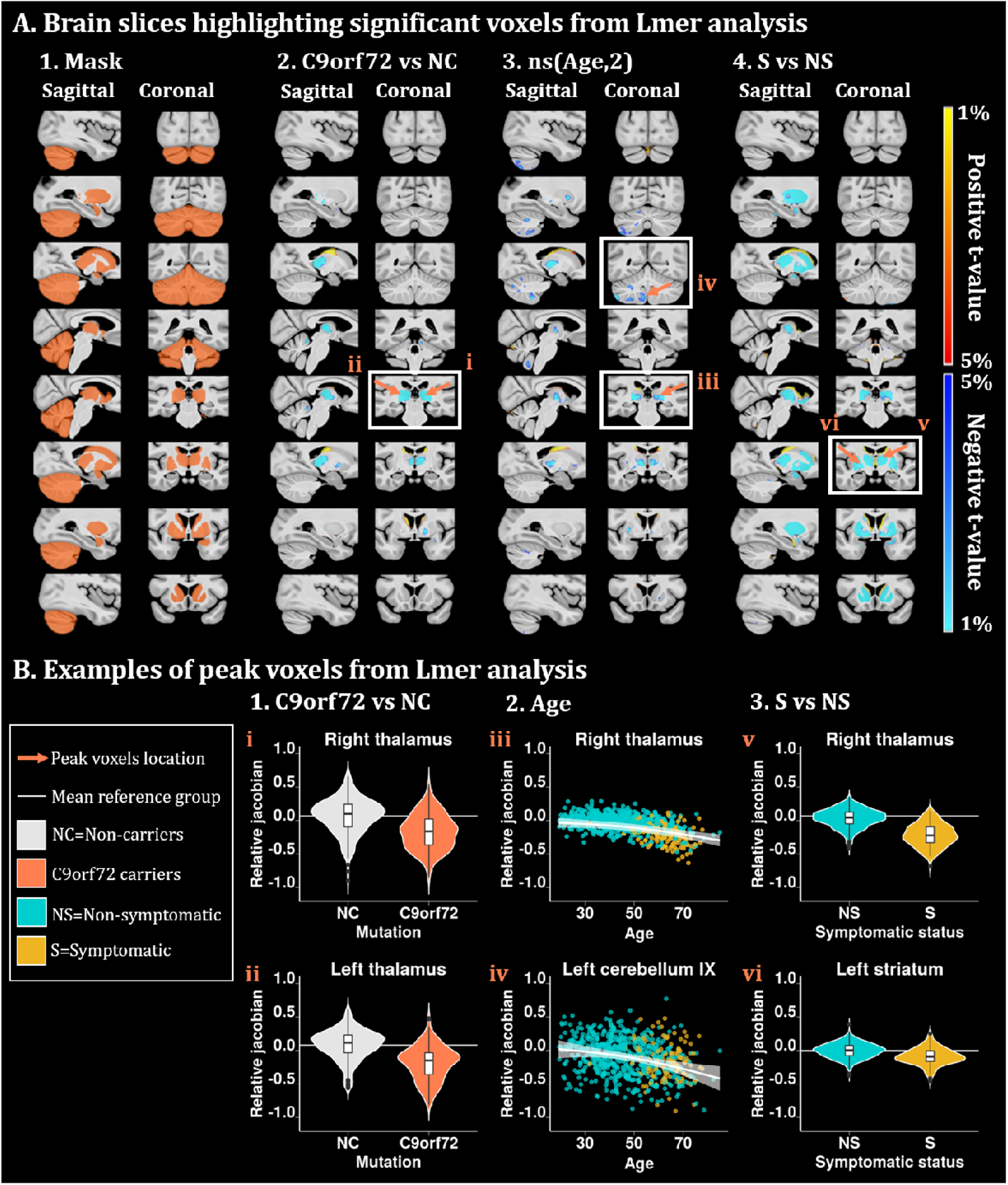
A. Brain slices of the brain highlighting significant voxels from the lmer analyses. t-value maps correspond to significant p-values between 5% and 1% after FDR correction. Axial slices represented from left to right and coronal slices represented from posterior to anterior. The t-statistics color maps for the significant expansion are in yellow to red and for the significant contraction are in turquoise to blue. White boxes and orange arrows were used to highlight the peak voxels selected for the plots in figure 2B. **A1**. Sagittal and coronal slices of the mask used to focus the analyses in the regions of interest. **A2**. Slices of the brain showing significant differences between the relative Jacobians of the *C9orf72* carriers versus the relative Jacobians of the non-carriers participants. **A3**. Slices of the brain map exhibiting significant second order volume decrease with age. **A4**. Brain map showing a significant volume reduction in the symptomatic participants compared to the non-symptomatic participants (non-carriers + presymptomatic). **B. Examples of peak voxels from lmer analyses**. White horizontal line highlights the mean relative Jacobian of the reference group (either non-carriers or presymptomatic). **B1**. Violin plots illustrate the relative Jacobians difference of two peak voxels in the right and left thalamus between the *C9orf72* carriers and the non-carriers individuals. **B2**. Best fit models showing the second order relationships between the relative Jacobians and age, using the predicted Jacobians between age 19 and 85 for a subject of mean EYO and unweighted averages over the levels of sex, genetic mutation and symptomatic status. These plots highlight a second order volume decrease with advanced age in two peak voxels of the right thalamus and left cerebellum lobule IX. **B3**. Violin plot of two peak voxels illustrating volume reduction in the right thalamus and left striatum of the symptomatic individuals compared to the non-symptomatic participants.

### 2.5. Flat map visualization

Matlab (*2014b*) and SUIT toolbox were used to display lmer and PLS results into a surface-based flatmap representation of the cerebellum ^50–53^. We used *suit_isolate_seg, suit_normalize_dartel* and *suit_reslice_dartel* functions to transform our template brain and our statistical maps into SUIT space. Then, we used *suit_map2surf* and *suit_plotflatmap* functions to obtain our statistical maps in a cerebellar flatmap representation. Simplified atlases of the resting-state networks and of the task processing were summarized from previous results ^54,55^.

## 3. Results

### 3.1. Demographic and clinical information

After motion quality control (see 2.3.2. Raw quality control), 130 scans were excluded (supplementary table 1). From 853 participants, 8 failed our preprocessing and 115 individuals were excluded due to missing CBI-R information. Table 1 describes the demographic and clinical information of the 730 individuals used in subsequent analyses for which the preprocessing and processing steps passed our quality control (QC). These participants included 184 *C9orf72* carriers, 185 *GRN* carriers, 80 *MAPT* carriers and 281 controls.

### 3.2. Linear mixed effect model

Figure 2.A demonstrates the brain areas showing significant effect with some of the demographic and clinical information of interest (genetic group, age and symptomatic status). EYO was not significant and is therefore not included in the figure. Figure 2.B shows representative significant peak voxels (highlighted in white boxes and orange arrows in the brain maps of Figure 2.A) to visualize several key regions of interest. In Figure 2.A1, the first two columns highlight the mask used for the Lmer analyses on the sagittal and coronal slices.

#### 3.2.1. Genetic group variable

Figure 2.A2 illustrates the brain areas showing significant volume differences between *C9orf72* carriers (including a majority of presymptomatic subjects) versus non-carrier participants. *C9orf72* carriers demonstrate significant volume decrease in the bilateral thalamus. No significant differences were found between *GRN* carriers and *MAPT* carriers versus non-carriers (meaning that age was the primary driver of volume change in this predominantly asymptomatic group). Figure 2.B1 illustrates two peak voxels in the right and left thalamus, demonstrating an approximate 20% volume decrease in the *C9orf72* carriers compared to non-carriers individuals.

#### 3.2.2. Age

In Figure 2.A3, the brain map exhibits the voxels demonstrating significant second order volume decrease with age, independently of mutation status. These voxels are mostly situated in the thalamus and in the cerebellum, particularly in lobule IX. Figure 2.B2 exhibits two peak voxels of the right thalamus and left cerebellum lobule IX, highlighting a second order volume decrease with advanced age. Interestingly, the pattern of decrease with age is seen for the entire age range and not only for the symptomatic participants (especially in the cerebellum). Also, larger relative Jacobian variability in the cerebellum was observed compared to the thalamus.

#### 3.2.3. Symptomatic status

In Figure 2.A4, the brain map shows a significant volume reduction in the bilateral thalamus, striatum (mainly accumbens nucleus, putamen and ventral caudate), globus pallidus and amygdala in the symptomatic participants (from all mutation groups) compared to the non-symptomatic participants (non-carriers + presymptomatic mutation carriers individuals). Of note, no volume difference was found between symptomatic and non-symptomatic participants in the cerebellum. Further, from the lmer results, no significant relationship was found between the relative Jacobians and EYO. Finally, figure 2.B3 illustrates the reduced volume of the symptomatic individuals, in two peak voxels of the right thalamus and left striatum, compared to the non-symptomatic participants.

### 3.3. PLS results

#### 3.3.1. Latent variables

PLS results demonstrate one significant LV per genetic group, except for the non-carriers PLS run which had no significant LVs. Figure 3 illustrates these LVs, for which we can see the brain score maps (A) and their corresponding behaviour scores (B) for each mutation group. Supplementary figure 1 demonstrates that even with a more lenient threshold (p<0.05), each genetic group appeared to still have a different pattern of cerebellar atrophy.

**Figure 3:**
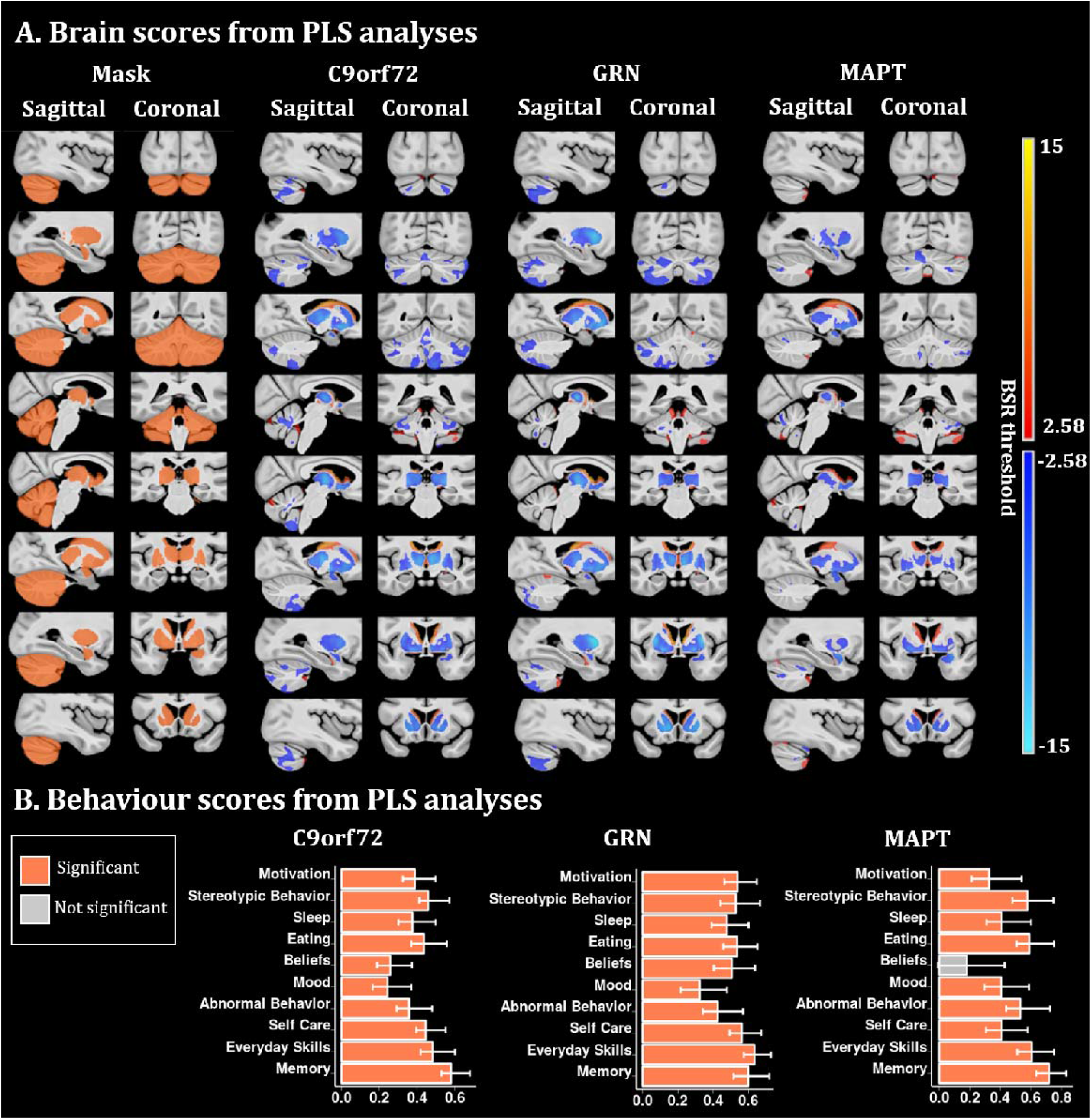
PLS analyses between the voxel-wise relative Jacobians and the CBI-R variables for each mutation group separately. **A)** Brain scores of each latent variable (LV) were plotted using the vertex wise BSR thresholded at 2.58 (p<0.01). The range of BSR values was [−12.4,11.7] for *C9orf72*, [−14.8,14.5] for *GRN* and [−8.4,9.1] for *MAPT* LV. A common minimum/maximum BSR threshold was selected [−15,15] to have a similar color scale between each brain map. Each group demonstrated one significant LV except the non-carriers group (not shown). The LV explained 91.8 % of the variance for *C9orf72*, 93.2 % of the variance for *GRN* and 84.4 % of the variance for *MAPT*. **B)** Bar plots describe the correlation of each CBI-R variable with each LV, with error bars denoting the 95% confidence interval. Orange color represents CBI-R variables that significantly participate in the LV while grey color represents non-significant CBI-R variables.

Overall, all the genetic groups demonstrated a volume reduction in the thalamus, striatum and globus pallidus being (to a lesser extent in the *MAPT* mutation group) associated with lower behaviour scores. The main difference between the maps was from the cerebellum, which displayed a large volume reduction in the *C9orf72* carriers, a moderate volume reduction in the *GRN* mutation groups and a much lesser volume reduction in the *MAPT* mutation group.

##### 3.3.1.1. C9orf72

*C9orf72* LV explained 91.8 % of the variance. *C9orf72* brain map demonstrates a pattern of volume reduction in the thalamus, globus pallidus, striatum and in a large area in the cerebellum, including superior posterior lobules, and lobules IX associated with higher behaviour scores (worse cognition and behaviour symptoms).

##### 3.3.1.2. GRN

*GRN* LV explained 93.2 % of the variance and the *GRN* map illustrates a volume reduction of the thalamus, globus pallidus, striatum and very subtle parts of the cerebellum, mostly in lobules Crus I and Crus II. The *GRN* behaviour scores plot demonstrates significantly worse behaviour scores being linked to a volume reduction in the structures described above.

##### 3.3.1.3. MAPT

Finally, the *MAPT* LV explained 84.4 % of the variance and its map demonstrates a volume reduction in the thalamus, globus pallidus, striatum, larger effects in the amygdala and very subtle effects in the cerebellum. The MAPT behaviour scores plot demonstrated significantly worse behaviour scores in all CBI-R categories, except for the beliefs.

#### 3.3.2. Post-hoc analyses of PLS results

Brain and behaviour scores from PLS analyses were analysed to examine if they could be explained by the demographic and clinical information of the participants.

Overall, *C9orf72* and *MAPT* brain scores were mostly related to EYO and expressed a second order relationship showing a steep brain score decline up to 20 years before symptom onset. However, *GRN* brain scores were not related to EYO but age. Additionally, symptomatic participants were showing worse brain and behaviour scores (except for the brain scores of *C9orf72* carriers) compared to the pre-symptomatic participants.

##### 3.3.2.1. C9orf72

Figure 4A demonstrates that *C9orf72* brain scores were significantly reduced with increased age and EYO (p=8.7×10^−8^ and p=9.2×10^−5^ respectively); however it was not significantly different between presymptomatic and symptomatic carriers. Age is not related to behaviour scores while increasing EYO and being symptomatic are linked to higher behaviour scores in all subcategories (p=3.9×10^−2^ and p=4.7×10^−22^ respectively).

**Figure 4:**
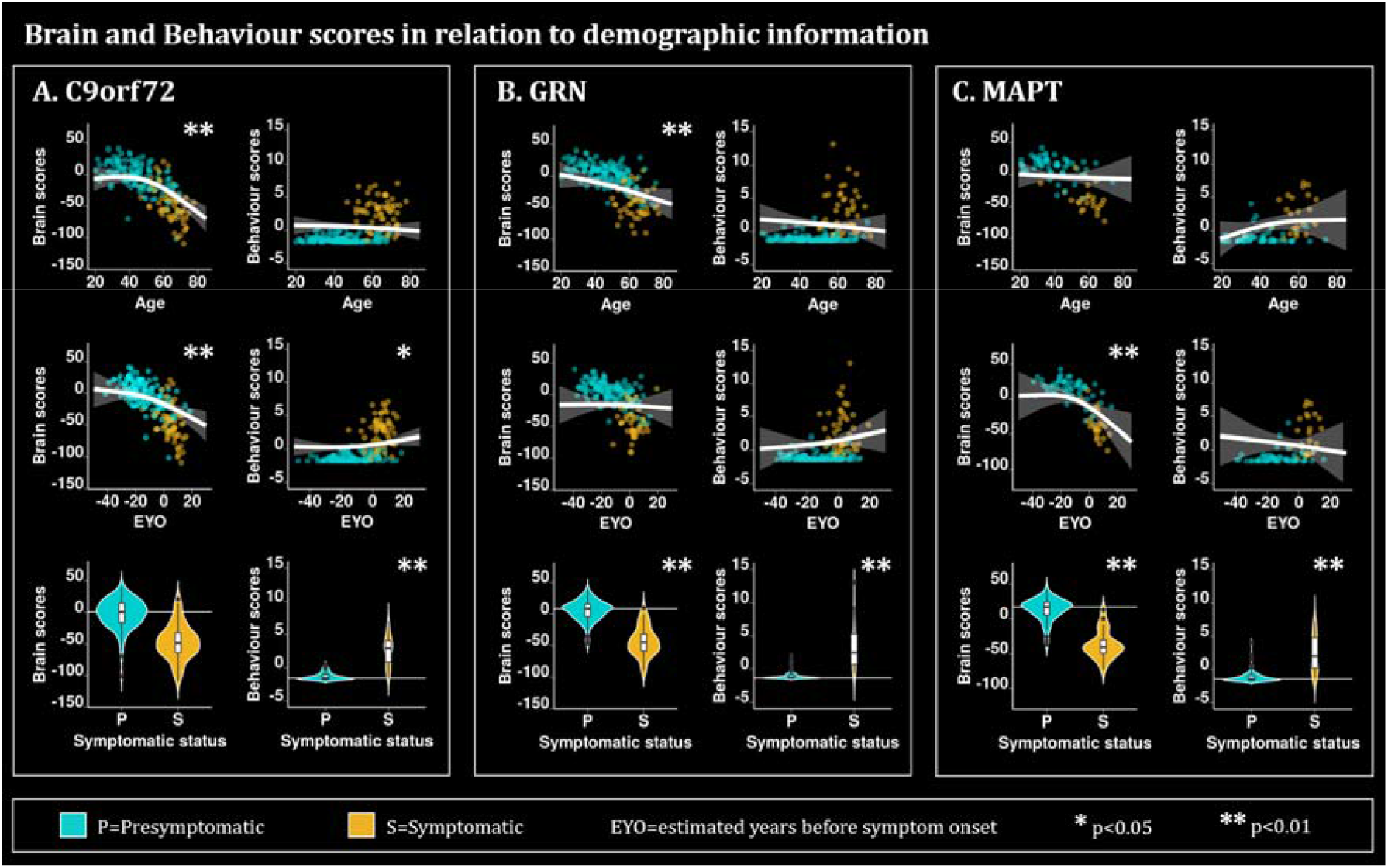
Plots describing the relationship of the brain and behaviour scores for **A**. *C9orf72*, **B**. *GRN* and **C**. *MAPT* mutation carriers with demographic and clinical information such as age, EYO, and symptomatic status. The plots for age and EYO either demonstrate the second order relationships between the relative Jacobians and age using the predicted Jacobians between age 19 and 85 for a subject of mean EYO or using the predicted Jacobians between EYO −50 and 30 for a subject of mean age, respectively. These models were computed using the unweighted averages over the levels of sex, education and symptomatic status. Turquoise is used to highlight the presymptomatic (P) individuals while gold is used to highlight the symptomatic (S) individuals. * is used to show significant variables (p<0.05 after FDR correction) and ** to show significant variables (p<0.01 after FDR correction. The age and EYO relationships were plotted based on the lmer model. White horizontal lines highlight the mean relative Jacobian of the presymptomatic individuals (reference group).

##### 3.3.2.2. GRN

In figure 4B, *GRN* brain scores are reduced with increasing age and being symptomatic (p=6.8×10^−3^ and 5.0×10^−16^ respectively), while EYO is not related to the brain scores. *GRN* behaviour scores are only related to being presymptomatic or symptomatic (p=2.6×10^−19^).

##### 3.3.2.3. MAPT

Finally, the brain scores of *MAPT* are significantly reduced with increased EYO and for symptomatic individuals (p=3.6×10^−3^ and p=6.0×10^−7^ respectively), while the behaviour scores are only increased for the symptomatic individuals compared to presymptomatic individuals (p=4.3×10^−5^). Interestingly, for both *C9orf72* and *MAPT*, the brain scores are starting to decrease up to 20 years before the expected symptom onset, when most of the individuals are still presymptomatic (turquoise color; also seen in supplementary figure 1 where only presymptomatic individuals were tested).

### 3.4. Cerebellar contribution

Figure 5 summarizes our main cerebellar results transformed into flat maps and compares them with simplified atlases of cerebellar anatomy (Figure 5A), resting-state network (Figure 5B) and task processing (Figure 5C), created based on previous findings ^50,54,55^. Figure 5D demonstrates the different brain maps extracted from PLS analyses for each mutation group. Of note, the *C9orf72* map is principally located on lobules Crus I, Crus II, VIIIB and IX. The *GRN* map is mainly localised in lobules Crus I and Crus II while the *MAPT* map only includes a very small part of lobule VI. Interestingly, we observe that the *C9orf72* map overlaps mostly with the default mode network, and corresponds with the working memory, language and social tasks. On the other hand, the *GRN* map overlaps mostly with the frontoparietal network and the working memory task.

**Figure 5:**
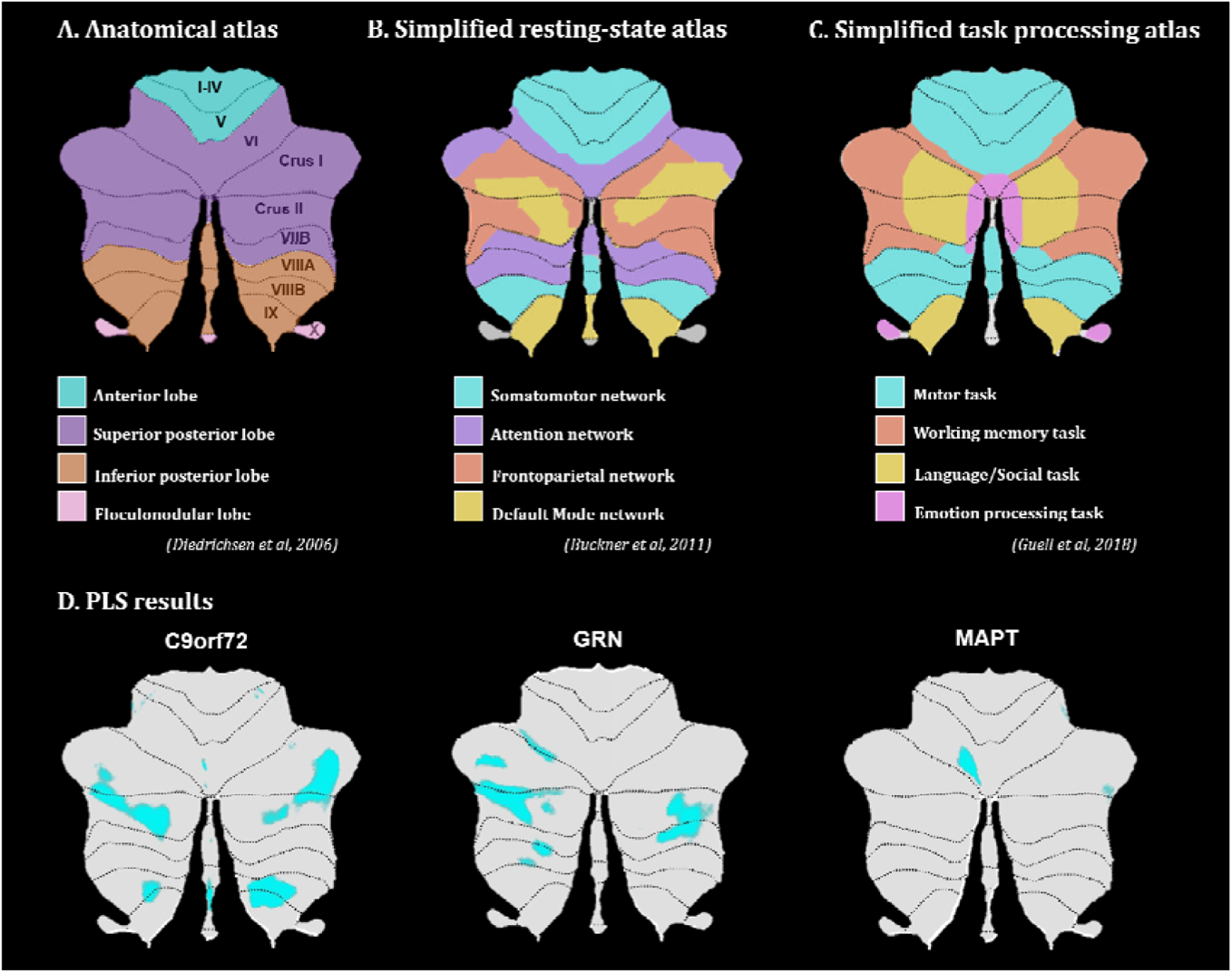
Flatmaps of the **A)** cerebellar anatomical atlas (Diedrichsen, 2006; Diedrichsen *et al*., 2009), **B)** simplified resting-state network atlas (Buckner *et al*., 2011), **C)** simplified task processing atlas (Guell, Gabrieli and Schmahmann, no date) and **D)** LV1 brain map results from PLS analyses for each mutation group.

## Discussion

The objective of this paper was to further our understanding of the role of subcortical brain atrophy in behavioural symptoms in the genetic forms of FTD. Here, we used DBM, univariate voxel-wise analyses and multivariate techniques to disentangle these relationships. First, all the individuals (mutation non-carriers and carriers) had progressively lower volume in the cerebellar lobule IX and in the thalamus throughout the adult lifespan. Second, when pooling at-risk and symptomatic subjects, we found a significantly smaller volume in the thalamus of *C9orf72* carriers compared to non-carriers. However, no significant differences were found between *GRN* or MAPT carriers (including presymptomatic and symptomatic) and non-carrier individuals. Third, we found that symptomatic individuals demonstrated smaller volumes in the thalamus, striatum, globus pallidus and amygdala compared to non-symptomatic individuals (either non-carriers or asymptomatic carriers). Overall, these univariate analyses demonstrate direct effects of age, C9orf72 expansion and symptomatic status on the subcortical volumes.

Our PLS analyses demonstrated overlapping association between brain and behaviour, but with specificity corresponding to each genetic group. While all brain maps included the thalamus, globus pallidus and striatum, *C9orf72* demonstrated a unique atrophy of a large area of the cerebellum including lobules Crus I, Crus II, VIIIB and IX, which is in agreement with a recent study showing preferential cerebellar involvement in *C9orf72* carriers ^56^. *GRN* cerebellum map demonstrated more restricted atrophy of the cerebellar lobules Crus I and Crus II. *MAPT* LV included a very limited atrophy of the cerebellum but a larger atrophy of the amygdala compared to the other two genetic groups. These findings are in line with previous studies showing thalamic and cerebellar symmetrical and widespread patterns of atrophy in *C9orf72* expansion carriers ^57,58^, while amygdalar atrophy seemed to be more related to *MAPT* carriers (in line with its predominant antero-medial temporal atrophy ^18,56^.

While we saw brain map differences between mutation groups, the PLS behavioural results did not demonstrate differentiated effects. Therefore the variations in atrophy observed across genetic groups did not lead to symptomatic profile differences. This is consistent with clinical observations which suggest that different mutations have major overlap in their clinical presentations, despite regional specificity in atrophy profiles ^9^. Additionally, our PLS analysis on the non-carrier group did not lead to a significant LV demonstrating that these patterns of brain/behaviour are specific to mutation carriers, as opposed to normative anatomical correlates of behavioural traits in healthy adults. Given the significant prevalence of psychotic symptoms in C9orf72 and to a lesser extent in GRN, we would have expected to find a specific latent variable for psychosis-related categories (beliefs or abnormal behaviour). However, the absence of such results could be explained by the relatively small number of symptomatic carriers with psychotic symptoms compared to the number of presymptomatic carriers. Of note, this lack of brain/behaviour differences between genetic groups, is in contrast to some previous reports that link specific symptoms to specific anatomical correlates ^24,59^. However, while these previous studies have found significant association between behaviour symptoms and genetic groups, they ran a series of ANOVA or a series of linear models for each symptom. On the contrary, here we used PLS analyses which is a recommended technique when the number of explanatory variables is high, and where it is likely that the explanatory variables are correlated ^45^, which is the case in our study.

As expected, the presence of symptoms was consistently a strong predictor of the PLS-derived brain and behaviour scores. However, different relationships of these scores with age and EYO were found for each genetic group. Indeed, *C9orf72* and *MAPT* brain scores were related to EYO and expressed a second order relationship highlighting a steeper brain score decline 20 years before symptom onset. However, *GRN* brain scores were related to age and not EYO. These results are consistent with previous reports that *GRN* mutation carrier symptom onset is only weakly predictable using parental age of onset, while *MAPT* and *C9orf72* EYO were more predictable ^27^. Also, a previous study has shown that while variant-specific grey matter atrophy (mostly in the thalamus) has been shown to start up to 20 years before EYO in *C9orf72* carriers, atrophy starts 15 years before EYO for *MAPT* carriers (mostly fronto-parietal) and only 10 years before EYO for GRN carriers (mostly medial-temporal) ^6^. Further, EYO was a better predictor at explaining the covariance between morphometry and CBI-R variables (at least for *C9orf72* and *MAPT* carriers), but it was not directly associated with morphometry (from our univariate analyses). These results suggest that future clinical and research studies might benefit from using brain atrophy and behaviour variables simultaneously to identify subjects at higher risk of short-term clinical conversion to the dementia stage. In addition, our results suggest that the brain regions that we found to be significantly associated with the CBI-R scores (subcortical structures + cerebellum Crus I and II, lobule VIIIB and IX) could be particularly sensitive to changes related to EYO rather than doing whole brain voxel-wise analyses to predict EYO.

Further, our results suggest that while we did find differences in the pattern of atrophy of the cerebellum and amygdala in each genetic group, we also found that the striatum, thalamus and globus pallidus were consistently impacted across the mutation carriers; in line with the strong overlap of clinical symptoms. While we identified a similar behavioural cluster in all the genetic groups, we can hypothesize that the different patterns of cerebellar and amygdalar atrophy could be responsible for symptomatic differences in each mutation group later in the disease progression. For example, this could contribute to the more pronounced psychotic symptoms (beliefs or abnormal behaviour) in *C9orf72* carriers (potentially linked to the preferential cerebellar lobule Crus I, Crus II and lobule IX atrophy) while *MAPT* carriers demonstrate more depression and anxiety (potentially linked to the preferential amygdalar atrophy).

Of interest, the cerebellar structures found to be associated with the CBI-R variables (Crus I, Crus II and IX) correspond to the ones previously found to be related with frontoparietal and default mode networks (DMN; ^60^). In our results, cerebellar atrophy in *C9orf72* carriers was predominantly related to the DMN, which relates to previous findings ^61^, while in *GRN* carriers, cerebellar regions were primarily functionally correlated to the fronto-parietal network, which ties in to the well documented component of parietal involvement in this disease ^18^. Interestingly, reduction of Crus I and Crus II volumes has previously been associated with behavioural and cognitive deficits in FTD ^62^. These results suggest that while our study focused on the anatomical changes in FTD in relation with behaviour, we demonstrated that the structures covarying the most with behavioural changes colocalize specifically with the DMN and fronto-parietal networks, motivating the relevance of using fMRI techniques to study brain/behaviour changes in familial FTD.

Some limitations in our study need to be highlighted and discussed. Indeed, one of the main conceivable biases in our results come from the limited number of *MAPT* mutation carriers in the GENFI dataset, which might explain the reduced brain involvement in this genetic group. Indeed, only about 18% of our total number of mutation carriers were *MAPT* carriers. However worldwide, *MAPT* mutation is also the least frequent (23.2%), followed by *GRN* mutation (34.6%) and *C9orf72* expansion (42.1%) ^27^. Therefore, our dataset was quite representative of the general mutation distribution in FTD. Further, while GENFI is the largest dataset of carriers for FTD-linked mutations to date and offers a sufficient number of participants to conduct complex statistical analyses, only 18% of those participants were symptomatic mutation carriers, preventing us from testing different age-relationships for symptomatic vs asymptomatic individuals based on our stringent statistical approach. Indeed, the AIC method selection values the goodness of fit as well as the simplicity of the model (trade-off between the risk of overfitting and the risk of underfitting) and in our dataset the interaction between age and mutation group was considered to be less good of a fit and was excluded from the model. While this suggests that the subcortical volumetric effect of GRN and MAPT in presymptomatic carriers was too small to have a meaningful contribution to the overall model fit, it does not invalidate previous findings relating those structures to late-presymptomatic and symptomatic disease stages.

Secondly, an important consideration is that this study aims to understand the brain and CBI-R variables impairment during the presymptomatic phase of the disease, and uses EYO as an estimator of age of symptom onset. However, as discussed above, EYO accuracy is far from perfect and varies between genetic groups ^27^. Therefore, in order to better characterize the evolution of the brain and behaviour during this presymptomatic phase, longitudinal data with documentation of the actual age at onset would be necessary. This will eventually be feasible with future GENFI date release since follow up brain imaging and CBI-R assessments are performed longitudinally on these individuals. Finally, it is important to note that while our results allowed straightforward qualitative comparisons between each PLS findings, no test has been performed to quantitatively compare the brain maps of each genetic group. Further, in order to verify that our differences in brain maps were not driven by a threshold effect, we also plotted the different brain maps using a less stringent significant threshold (Supplementary figure 1), demonstrating that even with a more lenient threshold, different parts of the cerebellum were associated with the behaviour scores for each genetic group.

To conclude, our study shows that the subcortical and cerebellar circuitry is heavily involved in the relationship between FTD pathology and the emergence of neuropsychiatric symptoms, even in the presymptomatic stage. Indeed, we have shown that *C9orf72* and *MAPT* brain scores start to decrease up to 20 years before the expected symptom onset, when most individuals are still presymptomatic. Furthermore, while all brain maps included the thalamus, globus pallidus and striatum, *C9orf72* demonstrated a unique involvement of a large area of the cerebellum, including Crus I and II, lobule VIIIB and IX, and *MAPT* brain pattern included a very limited part of the cerebellum but involved more of the amygdala compared to the other two genetic groups. Finally, the variations in atrophy observed across genetic groups did not explain symptomatic profile differences, suggesting that these differences in atrophy pattern in the cerebellum and amygdala might only lead to behavioural differences later in the disease progression.

## Abbreviations

AIC: Akaike information criterion
C9orf72: chromosome 9 open reading frame 72
CBI-R: Cambridge Behavioural Inventory Revised
CoBrA: Computational Brain Anatomy
DBM: deformation-based morphometry
EYO: estimated years before the age of symptom onset
FTD: frontotemporal dementia
GENFI: Genetic Frontotemporal dementia Initiative
GRN: progranulin
LMER: linear mixed effect models
LV: latent variable
MAPT: microtubule-associated protein tau
MNI: Montreal neurological institute
MRI: magnetic resonance imaging
PLS: partial least squares
QC: quality control
ROI: region of interest
SVD: singular value decomposition
BK-1: TANK-binding kinase 1

## Figure legends

**Figure 1: Chart flow of the step by step methods and analyses used in this paper. 1- Green**: Raw inputs (2.3.2. Raw quality control); **2-Gray**: preprocessing (2.3.3 Preprocessing); **3-Orange**: Deformation based morphometry (2.3.4. Deformation based morphometry); **4-Purple**: Mask creation (2.3.5. Mask creation); **5-Turquoise**: linear mixed effect models (2.4.1. Linear mixed effect models); **6-Pink**: Partial least squares (2.4.2. Partial least squares analysis). Abbreviations: QC: quality control; Preprocessing: minc-bpipe-library; DBM: deformation based morphometry; SVD: singular value decomposition; PLS: partial least squares; LMER: linear mixed effect models.

**Figure 2: A. Brain slices of the brain highlighting significant voxels from the lmer analyses**. t-value maps correspond to significant p-values between 5% and 1% after FDR correction. Axial slices represented from left to right and coronal slices represented from posterior to anterior. The t-statistics color maps for the significant expansion are in yellow to red and for the significant contraction are in turquoise to blue. White boxes and orange arrows were used to highlight the peak voxels selected for the plots in figure 2B. **A1**. Sagittal and coronal slices of the mask used to focus the analyses in the regions of interest. **A2**. Slices of the brain showing significant differences between the relative Jacobians of the *C9orf72* carriers versus the relative Jacobians of the non-carriers participants. **A3**. Slices of the brain map exhibiting significant second order volume decrease with age. **A4**. Brain map showing a significant volume reduction in the symptomatic participants compared to the non-symptomatic participants (non-carriers + presymptomatic). **B. Examples of peak voxels from lmer analyses**. White horizontal line highlights the mean relative Jacobian of the reference group (either non-carriers or presymptomatic). **B1**. Violin plots illustrate the relative Jacobians difference of two peak voxels in the right and left thalamus between the *C9orf72* carriers and the non-carriers individuals. **B2**. Best fit models showing the second order relationships between the relative Jacobians and age, using the predicted Jacobians between age 19 and 85 for a subject of mean EYO and unweighted averages over the levels of sex, genetic mutation and symptomatic status. These plots highlight a second order volume decrease with advanced age in two peak voxels of the right thalamus and left cerebellum lobule IX. **B3**. Violin plot of two peak voxels illustrating volume reduction in the right thalamus and left striatum of the symptomatic individuals compared to the non-symptomatic participants.

**Figure 3:** PLS analyses between the voxel-wise relative Jacobians and the CBI-R variables for each mutation group separately. **A)** Brain scores of each latent variable (LV) were plotted using the vertex wise BSR thresholded at 2.58 (p<0.01). The range of BSR values was [−12.4,11.7] for *C9orf72*, [−14.8,14.5] for *GRN* and [−8.4,9.1] for *MAPT* LV. A common minimum/maximum BSR threshold was selected [−15,15] to have a similar color scale between each brain map. Each group demonstrated one significant LV except the non-carriers group (not shown). The LV explained 91.8 % of the variance for *C9orf72*, 93.2 % of the variance for *GRN* and 84.4 % of the variance for *MAPT*. **B)** Bar plots describe the correlation of each CBI-R variable with each LV, with error bars denoting the 95% confidence interval. Orange color represents CBI-R variables that significantly participate in the LV while grey color represents non-significant CBI-R variables.

**Figure 4:** Plots describing the relationship of the brain and behaviour scores for **A**. *C9orf72, GRN* and **C**. *MAPT* mutation carriers with demographic and clinical information such as age, EYO, and symptomatic status. The plots for age and EYO either demonstrate the second order relationships between the relative Jacobians and age using the predicted Jacobians between age 19 and 85 for a subject of mean EYO or using the predicted Jacobians between EYO −50 and 30 for a subject of mean age, respectively. These models were computed using the unweighted averages over the levels of sex, education and symptomatic status. Turquoise is used to highlight the presymptomatic (P) individuals while gold is used to highlight the symptomatic (S) individuals. * is used to show significant variables (p<0.05 after FDR correction) and ** to show significant variables (p<0.01 after FDR correction. The age and EYO relationships were plotted based on the lmer model. White horizontal lines highlight the mean relative Jacobian of the presymptomatic individuals (reference group).

**Figure 5:** Flatmaps of the **A)** cerebellar anatomical atlas ^50,51^, **B)** simplified resting-state network atlas ^55^, **C)** simplified task processing atlas ^54^ and **D)** LV1 brain map results from PLS analyses for each mutation group.

## References

1. Coyle-Gilchrist, I. T. S. et al. Prevalence, characteristics, and survival of frontotemporal lobar degeneration syndromes. Neurology 86, 1736–1743 (2016).

2. Ratnavalli, E., Brayne, C., Dawson, K. & Hodges, J. R. The prevalence of frontotemporal dementia. Neurology 58, 1615–1621 (2002).

3. Mendez, M. F., Shapira, J. S., Woods, R. J., Licht, E. A. & Saul, R. E. Psychotic Symptoms in Frontotemporal Dementia: Prevalence and Review. Dementia and Geriatric Cognitive Disorders vol. 25 206–211 (2008).

4. Mourik, J. C. et al. Frontotemporal dementia: behavioral symptoms and caregiver distress. Dement. Geriatr. Cogn. Disord. 18, 299–306 (2004).

5. Takada, L. T. The Genetics of Monogenic Frontotemporal Dementia. Dement Neuropsychol 9, 219–229 (2015).

6. Greaves, C. V. & Rohrer, J. D. An update on genetic frontotemporal dementia. J. Neurol. 266, 2075–2086 (2019).

7. Rohrer, J. D. et al. The heritability and genetics of frontotemporal lobar degeneration. Neurology 73, 1451–1456 (2009).

8. Bertrand, A. et al. Early Cognitive, Structural, and Microstructural Changes in Presymptomatic C9orf72 Carriers Younger Than 40 Years. JAMA Neurol. 75, 236–245 (2018).

9. Rohrer, J. D. et al. Presymptomatic cognitive and neuroanatomical changes in genetic frontotemporal dementia in the Genetic Frontotemporal dementia Initiative (GENFI) study: a cross-sectional analysis. Lancet Neurol. 14, 253–262 (2015).

10. Le Blanc, G. et al. Faster Cortical Thinning and Surface Area Loss in Presymptomatic and Symptomatic C9orf72 Repeat Expansion Adult Carriers. Ann. Neurol. 88, 113–122 (2020).

11. Ducharme, S., Bajestan, S., Dickerson, B. C. & Voon, V. Psychiatric Presentations of C9orf72 Mutation: What Are the Diagnostic Implications for Clinicians? J. Neuropsychiatry Clin. Neurosci. 29, 195–205 (2017).

12. Tavares, T. P. et al. Early symptoms in symptomatic and preclinical genetic frontotemporal lobar degeneration. J. Neurol. Neurosurg. Psychiatry 91, 975–984 (2020).

13. Du, A.-T. et al. Different regional patterns of cortical thinning in Alzheimer’s disease and frontotemporal dementia. Brain 130, 1159–1166 (2007).

14. Rohrer, J. D. et al. Patterns of cortical thinning in the language variants of frontotemporal lobar degeneration. Neurology 72, 1562–1569 (2009).

15. Hartikainen, P. et al. Cortical thickness in frontotemporal dementia, mild cognitive impairment, and Alzheimer’s disease. J. Alzheimers. Dis. 30, 857–874 (2012).

16. Borrego-Écija, S. et al. Disease-related cortical thinning in presymptomatic granulin mutation carriers. Neuroimage Clin 29, 102540 (2021).

17. Ratti, E. et al. Regional prefrontal cortical atrophy predicts specific cognitive-behavioral symptoms in ALS-FTD. Brain Imaging and Behavior (2021) doi:10.1007/s11682-021-00456-1.

18. Cash, D. M. et al. Patterns of gray matter atrophy in genetic frontotemporal dementia: results from the GENFI study. Neurobiol. Aging 62, 191–196 (2018).

19. Sherman, S. M. Thalamus plays a central role in ongoing cortical functioning. Nat. Neurosci. 19, 533–541 (2016).

20. Balleine, B. W., Delgado, M. R. & Hikosaka, O. The role of the dorsal striatum in reward and decision-making. J. Neurosci. 27, 8161–8165 (2007).

21. Gil Robles, S., Gatignol, P., Capelle, L., Mitchell, M.-C. & Duffau, H. The role of dominant striatum in language: a study using intraoperative electrical stimulations. J. Neurol. Neurosurg. Psychiatry 76, 940–946 (2005).

22. Cardinal, R. N., Parkinson, J. A., Hall, J. & Everitt, B. J. Emotion and motivation: the role of the amygdala, ventral striatum, and prefrontal cortex. Neurosci. Biobehav. Rev. 26, 321–352 (2002).

23. Schmahmann, J. D. & Caplan, D. Cognition, emotion and the cerebellum. Brain: a journal of neurology vol. 129 290–292 (2006).

24. Sellami, L. et al. Distinct Neuroanatomical Correlates of Neuropsychiatric Symptoms in the Three Main Forms of Genetic Frontotemporal Dementia in the GENFI Cohort. J. Alzheimers. Dis. 65, 147–163 (2018).

25. Krueger, C. E. et al. Longitudinal rates of lobar atrophy in frontotemporal dementia, semantic dementia, and Alzheimer’s disease. Alzheimer Dis. Assoc. Disord. 24, 43–48 (2010).

26. Devenney, E. M. et al. The neural correlates and clinical characteristics of psychosis in the frontotemporal dementia continuum and the C9orf72 expansion. Neuroimage Clin 13, 439–445 (2017).

27. Moore, K. M. et al. Age at symptom onset and death and disease duration in genetic frontotemporal dementia: an international retrospective cohort study. Lancet Neurol. 19, 145–156 (2020).

28. Wear, H. J. et al. The Cambridge Behavioural Inventory revised. Dement Neuropsychol 2, 102–107 (2008).

29. Bellon, E. M. et al. MR artifacts: a review. AJR Am. J. Roentgenol. 147, 1271–1281 (1986).

30. Smith, T. B. & Nayak, K. S. MRI artifacts and correction strategies. Imaging Med. (2010).

31. Reuter, M. et al. Head motion during MRI acquisition reduces gray matter volume and thickness estimates. Neuroimage 107, 107–115 (2015).

32. Bedford, S. A. et al. Large-scale analyses of the relationship between sex, age and intelligence quotient heterogeneity and cortical morphometry in autism spectrum disorder. Mol. Psychiatry 25, 614–628 (2020).

33. Tustison, N. J. et al. N4ITK: improved N3 bias correction. IEEE Trans. Med. Imaging 29, 1310–1320 (2010).

34. Collins, D. L., Neelin, P., Peters, T. M. & Evans, A. C. Automatic 3D intersubject registration of MR volumetric data in standardized Talairach space. J. Comput. Assist. Tomogr. 18, 192–205 (1994).

35. Dadar, M., Fonov, V. S., Collins, D. L. & Alzheimer’s Disease Neuroimaging Initiative. A comparison of publicly available linear MRI stereotaxic registration techniques. Neuroimage 174, 191–200 (2018).

36. Eskildsen, S. F. et al. BEaST: brain extraction based on nonlocal segmentation technique. Neuroimage 59, 2362–2373 (2012).

37. Avants, B. B. et al. A reproducible evaluation of ANTs similarity metric performance in brain image registration. Neuroimage 54, 2033–2044 (2011).

38. Akaike, H. A New Look at the Statistical Model Identification. Springer Series in Statistics 215–222 (1974) doi:10.1007/978-1-4612-1694-0_16.

39. Mazerolle, M. Improving data analysis in herpetology: using Akaike’s Information Criterion (AIC) to assess the strength of biological hypotheses. Amphib-reptil. 27, 169–180 (2006).

40. Tullo, S. et al. MR-based age-related effects on the striatum, globus pallidus, and thalamus in healthy individuals across the adult lifespan. Hum. Brain Mapp. 40, 5269–5288 (2019).

41. Bussy, A. et al. Hippocampal subfield volumes across the healthy lifespan and the effects of MR sequence on estimates. Cold Spring Harbor Laboratory 2020.05.28.121343 (2020) doi:10.1101/2020.05.28.121343.

42. Benjamini, Y. & Hochberg, Y. Controlling the False Discovery Rate: A Practical and Powerful Approach to Multiple Testing. J. R. Stat. Soc. Series B Stat. Methodol. 57, 289–300 (1995).

43. Benjamini, Y., Drai, D., Elmer, G., Kafkafi, N. & Golani, I. Controlling the false discovery rate in behavior genetics research. Behavioural brain research vol. 125 279–284 (2001).

44. McIntosh, A. R. & Lobaugh, N. J. Partial least squares analysis of neuroimaging data: applications and advances. Neuroimage 23 Suppl 1, S250–63 (2004).

45. McIntosh, A. R. & Mišić, B. Multivariate statistical analyses for neuroimaging data. Annu. Rev. Psychol. 64, 499–525 (2013).

46. Krishnan, A., Williams, L. J., McIntosh, A. R. & Abdi, H. Partial Least Squares (PLS) methods for neuroimaging: a tutorial and review. Neuroimage 56, 455–475 (2011).

47. Zeighami, Y. et al. A clinical-anatomical signature of Parkinson’s disease identified with partial least squares and magnetic resonance imaging. Neuroimage (2017) doi:10.1016/j.neuroimage.2017.12.050.

48. Patel, R. et al. Investigating microstructural variation in the human hippocampus using non-negative matrix factorization. Neuroimage 207, 116348 (2020).

49. Bussy, A. et al. Hippocampus shape across the healthy lifespan and its relationship with cognition. bioRxiv 2020.10.30.362921 (2020) doi:10.1101/2020.10.30.362921.

50. Diedrichsen, J. A spatially unbiased atlas template of the human cerebellum. Neuroimage 33, 127–138 (2006).

51. Diedrichsen, J., Balsters, J. H., Flavell, J., Cussans, E. & Ramnani, N. A probabilistic MR atlas of the human cerebellum. Neuroimage 46, 39–46 (2009).

52. Diedrichsen, J. et al. Imaging the deep cerebellar nuclei: a probabilistic atlas and normalization procedure. Neuroimage 54, 1786–1794 (2011).

53. Diedrichsen, J. & Zotow, E. Surface-Based Display of Volume-Averaged Cerebellar Imaging Data. PLoS One 10, e0133402 (2015).

54. Guell, X., Gabrieli, J. D. E. & Schmahmann, J. D. Triple Representation of Language, Working Memory, Social and Emotion Processing in the Cerebellum: Convergent Evidence from Task and Seed-Based Resting-State Fmri Analyses in a Single Large Cohort. doi:10.1101/254110.

55. Buckner, R. L., Krienen, F. M., Castellanos, A., Diaz, J. C. & Thomas Yeo, B. T. The organization of the human cerebellum estimated by intrinsic functional connectivity. Journal of Neurophysiology vol. 106 2322–2345 (2011).

56. Bocchetta, M. et al. Differential early subcortical involvement in genetic FTD within the GENFI cohort. Neuroimage Clin 30, 102646 (2021).

57. Sha, S. J. et al. Frontotemporal dementia due to C9ORF72 mutations: Clinical and imaging features. Neurology vol. 79 1002–1011 (2012).

58. Bocchetta, M. et al. Patterns of regional cerebellar atrophy in genetic frontotemporal dementia. Neuroimage Clin 11, 287–290 (2016).

59. Benussi, A. et al. Progression of Behavioral Disturbances and Neuropsychiatric Symptoms in Patients With Genetic Frontotemporal Dementia. JAMA Netw Open 4, e2030194 (2021).

60. Guell, X., Schmahmann, J. D., Gabrieli, J. D. E. & Ghosh, S. S. Functional gradients of the cerebellum. Elife 7, (2018).

61. Lee, S. E. et al. Altered network connectivity in frontotemporal dementia with C9orf72 hexanucleotide repeat expansion. Brain 137, 3047–3060 (2014).

62. Chen, Y., Kumfor, F., Landin-Romero, R., Irish, M. & Piguet, O. The Cerebellum in Frontotemporal Dementia: a Meta-Analysis of Neuroimaging Studies. Neuropsychol. Rev. 29, 450–464 (2019).

