## Supplementary material for "Cerebellar and subcortical atrophy contribute to psychiatric symptoms in frontotemporal dementia"

**Supplementary methods**

**Motion QC exclusion**

The 130 scans excluded were T1w images of 109 participants (some of them had several scans). The demographics of these excluded participants are given in Supplementary table 1. Most of the scans were from symptomatic participants (52.3%), while a similar number of non-carriers and presymptomatic carriers were excluded (24.8% and 22.9%, respectively).

**Supplementary table 1 :** Demographics of the participants excluded due to motion artifacts.

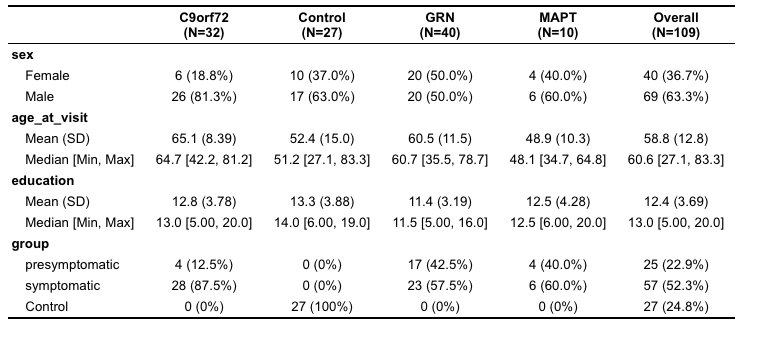

**Partial least squares analysis**

Partial least squares (PLS) correlation is a multivariate technique that is used to study and quantify the strength of the relationship between two matrices (Figure 1). The goal of PLS is to identify a set of latent variables (LVs) that explain patterns of covariance between “brain” and “CBI-R variables” data with the constraint that LVs explain as much of the covariance between the two matrices as possible. Theoretically, each LV depicts a linear combination of the “brain” and “CBI-R variables”. Here, three PLS analyses were run for each mutation group (C9orf72, GRN, MAPT and non-carriers separately). Our “brain” data included the relative Jacobian of each voxel for each subject (matrix size 3609356x184 for C9orf72; 3609356x185 for GRN, 3609356x80 for MAPT and 3609356x281 for non-carriers). Our “CBI-R variables” data contained individual and global CBI-R scores for each subject (matrix size 10x184 for C9orf72; 10x185 for GRN, 10x80 for MAPT and 10x281 for non-carriers). Note that this matrix does not contain any information on mutation carriers vs non carriers, symptomatic vs presymptomatic, age or EYO.

Each LV was tested statistically using permutation testing following a similar protocol as in previous studies (Anthony Randal McIntosh and Lobaugh 2004; Anthony R. McIntosh and Mišić 2013; Krishnan et al. 2011; Zeighami et al. 2017; Patel et al. 2020; Bussy et al. 2020). The rows of the input brain matrix were first permuted 10,000 times and PLS was performed on each new input brain matrix in order to obtain a distribution of singular values. From these testing, a nonparametric p-value was calculated for each LV to describe the probability that a singular value obtained from the original matrices was associated with chance. In order to highlight which LV had at least a 95% chance of not being associated with a random correlation in the original matrices, a p-value threshold of 0.05 was selected.

Secondly, now that some significant LVs were found, it is necessary to know the degree to which each brain and behaviour contributes to these LVs. Therefore, a bootstrap resampling technique was used. This time, both the rows of the brain and behaviour matrices were randomly sampled with replacement to generate 10,000 new sets of matrices with the goal of maintaining initial brain/behaviour relationships (unlike for the permutation testing described above). Then, each new matrix was subject to PLS in order to create a distribution of singular vector weights for each variable. A bootstrap ratio (BSR) representing the ratio of the singular vector weight over the standard error of the weight was calculated to assess the contribution and reliability of a brain variable. In this study, we used a BSR threshold of 2.58, which is analogous to a p-value of 0.01. Finally, a 95% confidence interval is obtained for each cognitive variable from the distribution of the singular vector weights (Krishnan et al., 2011; McIntosh and Lobaugh, 2004; Nordin et al., 2018; Persson et al., 2014; Zeighami et al., 2017).

**Visualization lmer model - code example**

model_brainscore <-lmer(GRN_Brain ~ ns(age_at_visit,2) + ns(eyo,2) + genetic_status_1 + education + sex +(1|scanner) + (1|blinded_family) , data=data_merged, control=lmerControl(optimizer="Nelder_Mead"))

fit_model_brainscore = as.data.frame(Effect(c( "eyo"), model_brainscore, xlevels=list(eyo=seq(-50,30,1)), given.values="equal"))

ggplot(data = data_merged, aes(y=GRN_Brain,x=eyo)) +

geom_point(aes(colour = factor(data_merged$genetic_status_1)), lwd=2,alpha=0.5,show.legend = FALSE) + scale_color_manual(values=c("cyan3", "goldenrod2"))+

geom_line(data = fit_model_brainscore, aes(y=fit),lwd=2, color="white") +

geom_ribbon(data = fit_model_brainscore, aes(y=fit, ymin=lower, ymax=upper), alpha=0.3, fill="white") +

theme_black()+ ylim(c(-140,70))+

theme(panel.grid.major = element_blank(), panel.grid.minor = element_blank())+

theme(legend.title=element_blank(), legend.position="right",legend.key.size = unit(0.5, "cm"))+

labs(x= "EYO", y = "Brain scores")+

theme(plot.title = element_text(size = 18, face = "bold"), legend.title=element_text(size=16), legend.text=element_text(size=16)) +

theme(axis.text.x = element_text(color = "white", size = 16, angle = 0, hjust = .5, vjust = .5, face = "bold"), axis.text.y = element_text(color = "white", size = 16, angle = 0, hjust = 1, vjust = 0, face = "bold"),

axis.title.x = element_text(color = "white", size = 16, angle = 0, hjust = .5, vjust = 0, face = "bold"),

axis.title.y = element_text(color = "white", size = 16, angle = 90, hjust = .5, vjust = .5, face = "bold"))+

theme(plot.margin = margin(0.6,.6,.6,.6, "cm"))

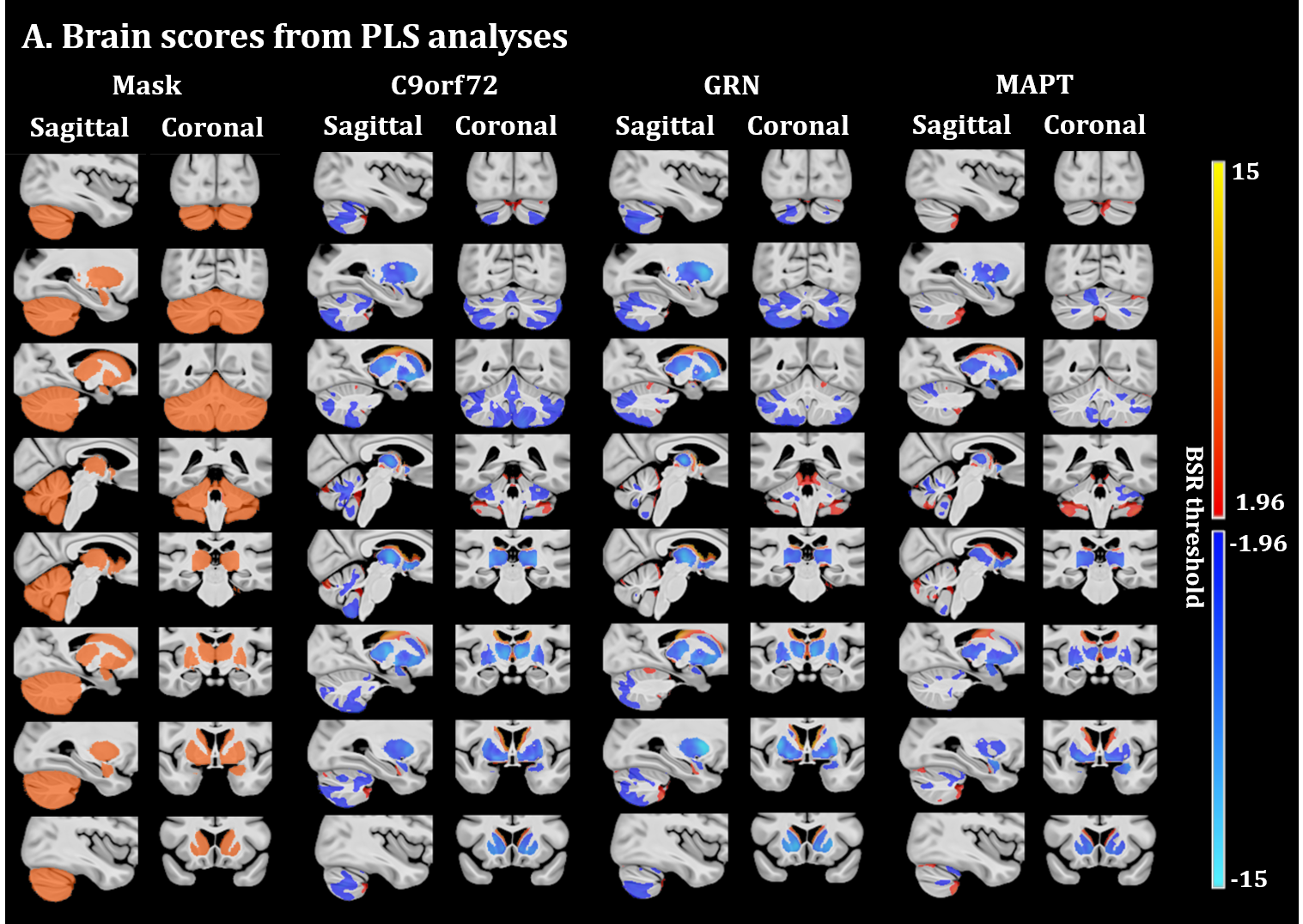

**Supplementary figure 1:** PLS analyses between the voxel-wise relative jacobians and the CBI-R variables for each mutation group separately. **A)** Brain scores of each latent variable (LV) were plotted using the vertex wise BSR thresholded at 1.96 (p<0.05). The range of BSR values was [-12.4,11.7] for *C9orf72*, [-14.8,14.5] for *GRN* and [-8.4,9.1] for *MAPT* LV. A common minimum/maximum BSR threshold was selected [-15,15] to have a similar color scale between each brain map. Each group demonstrated one significant LV except the non-carriers group (not shown). The LV explained 91.8 % of the variance for *C9orf72*, 93.2 % of the variance for *GRN* and 84.4 % of the variance for *MAPT*. **B)** Bar plots describe the correlation of each CBI-R variable with each LV, with error bars denoting the 95% confidence interval. Orange color represents CBI-R variables that significantly participate in the LV while grey color represents non-significant CBI-R variables.

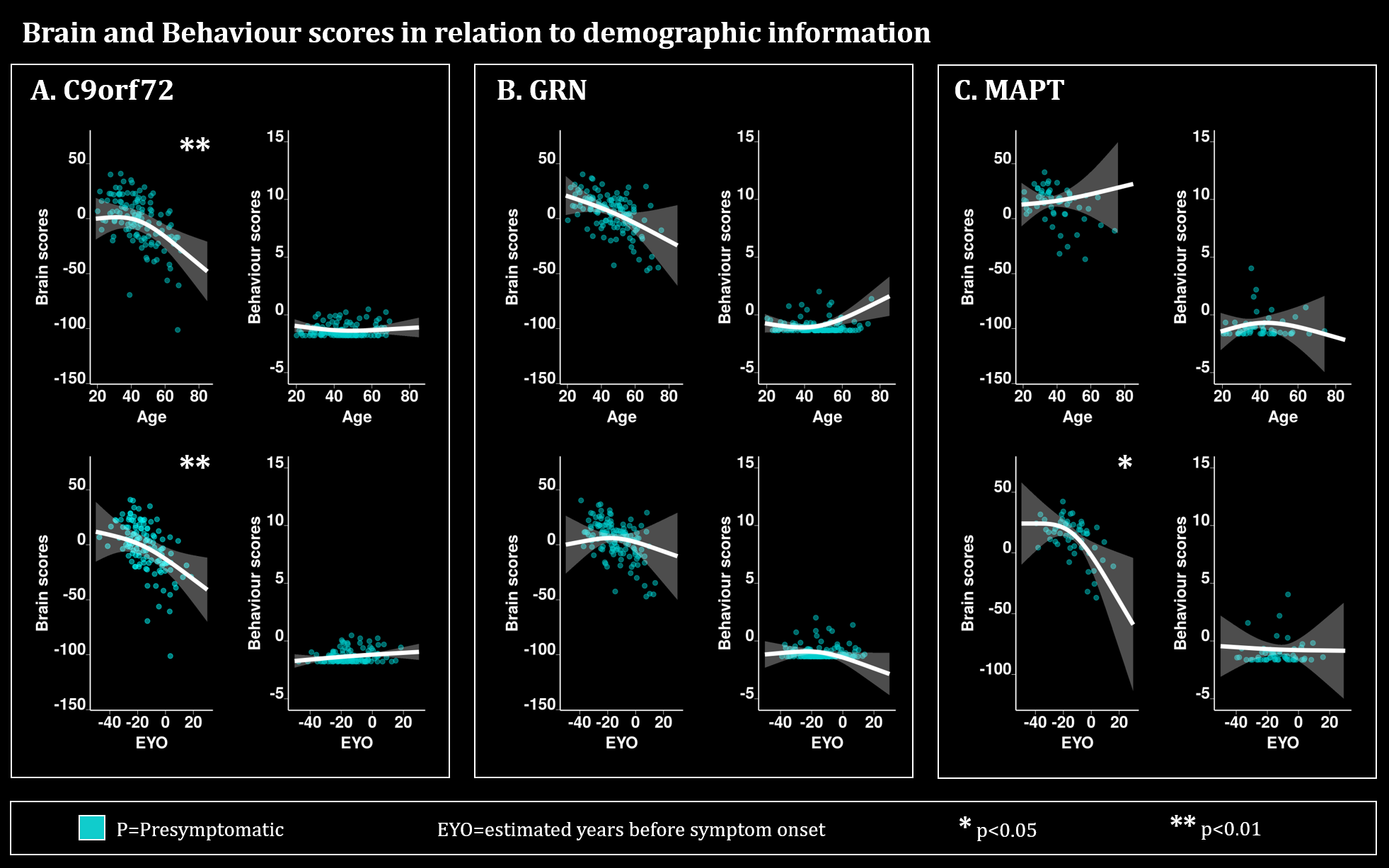

**Supplementary figure 2:** Plots describing the relationship of the brain and behaviour scores for the presymptomatic **A.** *C9orf72*, **B**. *GRN* and **C.** *MAPT* carrier individuals with demographic and clinical information such as age and EYO. The plots for age and EYO either demonstrate the second order relationships between the relative Jacobians and age using the predicted Jacobians between age 19 and 85 for a subject of mean EYO or using the predicted Jacobians between EYO -50 and 30 for a subject of mean age, respectively. These models were computed using the unweighted averages over the levels of sex, education and symptomatic status. Turquoise is used to highlight the presymptomatic (P) individuals, * is used to show significant variables (p<0.05 after FDR correction) and ** to show significant variables (p<0.01 after FDR correction. The age and EYO relationships were plotted based on the lmer model.
